# Modulation of prey size reveals adaptability and robustness in the cell cycle of an intracellular predator

**DOI:** 10.1101/2022.12.18.520956

**Authors:** Yoann G. Santin, Thomas Lamot, Jovana Kaljević, Renske van Raaphorst, Géraldine Laloux

## Abstract

Despite the remarkable diversity of bacterial lifestyles, the sophisticated regulatory networks underlying bacterial replication have only been investigated in a limited number of model species so far. In bacteria that do not rely on canonical binary division for proliferation, the coordination of major cellular processes is still mysterious. Moreover, bacterial growth and division remain largely unexplored within spatially confined niches where nutrients are limited. This includes the lifecycle of the model endobiotic predatory bacterium *Bdellovibrio bacteriovorus*, which grows by filamentation within its host or prey cells and produces a variable number of daughter cells. Here, we examined how the size of the micro-compartment in which predators replicate (i.e., the prey bacterium) impacts their cell cycle progression at the single-cell level. Using *Escherichia coli* with genetically encoded size differences, we show that the duration of the predator cell cycle scales with prey size. As a result, the size of the prey determines the number of predator offspring, through a relationship that is maintained across prey species. Strikingly, the nutritional quality of the prey cell determined the specific growth rate of predators regardless of prey size, reminiscent of the effect of medium composition on the growth of other bacteria. Tuning the predatory cell cycle by modulating prey dimensions also allowed us to reveal invariable temporal connections between key cellular processes. Altogether, our data uncover adaptability and robustness shaping the enclosed replication of *B. bacteriovorus*. Consequently, predators optimally exploit the finite prey resources and space while ensuring a strict cell cycle progression. This study opens the way to explore diverse cell cycle control strategies by extending their characterization to intracellular lifestyles, beyond canonical models and growth conditions.

## Introduction

Bacterial proliferation has been extensively studied for the past decades in a few model species (e.g., *Escherichia coli, Caulobacter crescentus* and *Bacillus subtilis*), revealing not only specific strategies but also general principles that govern streamlined cell cycle progression. For instance, these bacteria use exquisite mechanisms to tightly coordinate key cellular processes in space and time, such as cell growth, DNA replication, chromosome segregation, and cell division [1,2]. Examining how external cues impact the proliferation of those species further allowed to identify dependencies between fluctuating nutrients, cell size, and growth rate, which define the bacterial growth law [3–7]. However, little is known about the coordination of cell cycle events in non-model species and how environmental changes modulate their basic cellular processes. As the astonishing breadth of lifestyles in the bacterial realm is becoming increasingly appreciated, it is clear that the textbook picture of bacterial proliferation is not universal. For instance, several bacterial species are known to execute different types of division, some even producing more than two daughter cells per generation [8,9]. Furthermore, many species thrive in confined micro-environments where resources available to fuel their elongation and multiplication are finite – a limitation that has severely escaped investigation of bacterial growth so far. This is the case for intracellular pathogens that proliferate within eukaryotic host cells [10] or endobiotic predators that replicate inside other bacteria [11].

The model for endobiotic bacterial predation is *Bdellovibrio bacteriovorus*, which replicates within the periplasmic space of other diderm bacteria [12,13]. The cell cycle of *B. bacteriovorus* consists of two temporally separate phases. In the first so-called “attack” phase, motile and nonreproductive predators hunt for prey. Upon entry into the prey periplasm, *B. bacteriovorus* starts the second “growth” phase by digesting and feeding on the cellular content of the prey [14]. The predator cell then elongates in the bdelloplasts (i.e., the infected bacterium), forming a long filament before releasing variable numbers of progeny through multiple concomitant division events [15,16]. Recently, we used endogenous markers to show that *B. bacteriovorus* cells initiate asynchronous rounds of DNA replication, with multiple replisomes simultaneously active during filamentous growth, explaining how they achieve odd or even chromosome copy numbers [17]. In addition, the progressive segregation of the newly synthesized sister chromosomes results in a uniform distribution of the genetic material along the mother cell, allowing daughter cells to inherit precisely one chromosome upon division [17]. However, other key parameters underlying *B. bacteriovorus* cell cycle progression, such as the dynamics of filamentous growth and cell division, have never been studied quantitatively at the single-cell level [13]. Moreover, despite the fascinating obligate endobiotic lifestyle of *B. bacteriovorus*, it is still unclear whether the prey dimensions and content impact the predator cell cycle. It is generally assumed that larger prey cells release more predator daughter cells [15,18]. While this notion raises the intriguing possibility that the prey might shape the predator proliferative phase, the link between prey cell size and the number of predator offspring awaits unambiguous demonstration.

Here we used *B. bacteriovorus* as a model to systematically explore the impact of the size of the prey, i.e., the size of its closed replicative nest, on a non-canonical bacterial cell cycle. We followed markers of predator cell cycle progression by live fluorescence microscopy and developed automated tools to measure how the prey influences the predator cell cycle at the single-cell level. Differently sized prey strains allowed us to demonstrate that the timing of key cellular processes scales with prey size, suggesting that *B. bacteriovorus* adjusts its cell cycle to exploit its prey optimally. Furthermore, our data reveal the strict temporal coordination between predator chromosome dynamics, cell growth, and cell division, which is unperturbed by tuning the cell cycle duration. Finally, we assessed the impact of the prey’s nutritional quality on the specific growth rate of *B. bacteriovorus*. Altogether, our study provides the first quantitative and single-cell insights into the intracellular growth cycle of bacterial predators and its modulation by prey features.

## Results

### The number of predator offspring scales with prey size

To systematically test the effect of prey size on predator proliferation, we first asked whether varying prey cell dimensions impacts the number of newborn predators. We took advantage of previously described *E. coli* strains carrying point mutations on the actin homolog MreB (MreB*) in an otherwise identical genetic background, leading to different cell shapes and size distributions without altering cell physiology [19]. We selected the wild-type MG1655 and 3 MreB*\** strains (*ΔmreB* carrying an ectoptic copy of *mreB*_*WT*_, *mreB*_L251R_ or *mreB*_R193C_) as prey size variants, characterized by similar cell lengths but increasing widths. Hence, these four *E. coli* strains cover a broad range of cell sizes while limiting the variables to one dimension (**S1A Fig**). Since newborn *B. bacteriovorus* cells are monoploid [17], the number of chromosomes observed before predator division corresponds to the number of progeny. To exploit this relationship, each *E. coli* size variant was exposed to synchronized infection by a *B. bacteriovorus* strain natively producing the mCherry-tagged ParB protein (**Fig 1A**), which labels the chromosomal origin (*ori*) during the replication phase [17]. ParB-mCherry fluorescent foci were automatically detected from time-lapse fluorescence images, and the corresponding prey size was measured as the area of each infected cell (bdelloplast) at the start of the time-lapse (see Methods and **S2 Fig**). Our data show that the number of predator offspring tends to rise in prey strains of increasing size (**Fig 1A-B**). This trend was confirmed when preys were binned per area (**Fig 1C**) or at the single-cell level (**Fig 1D**) across all *E. coli* strains, indicating that this effect is independent of the *mreB* backgrounds.

**Fig 1.**
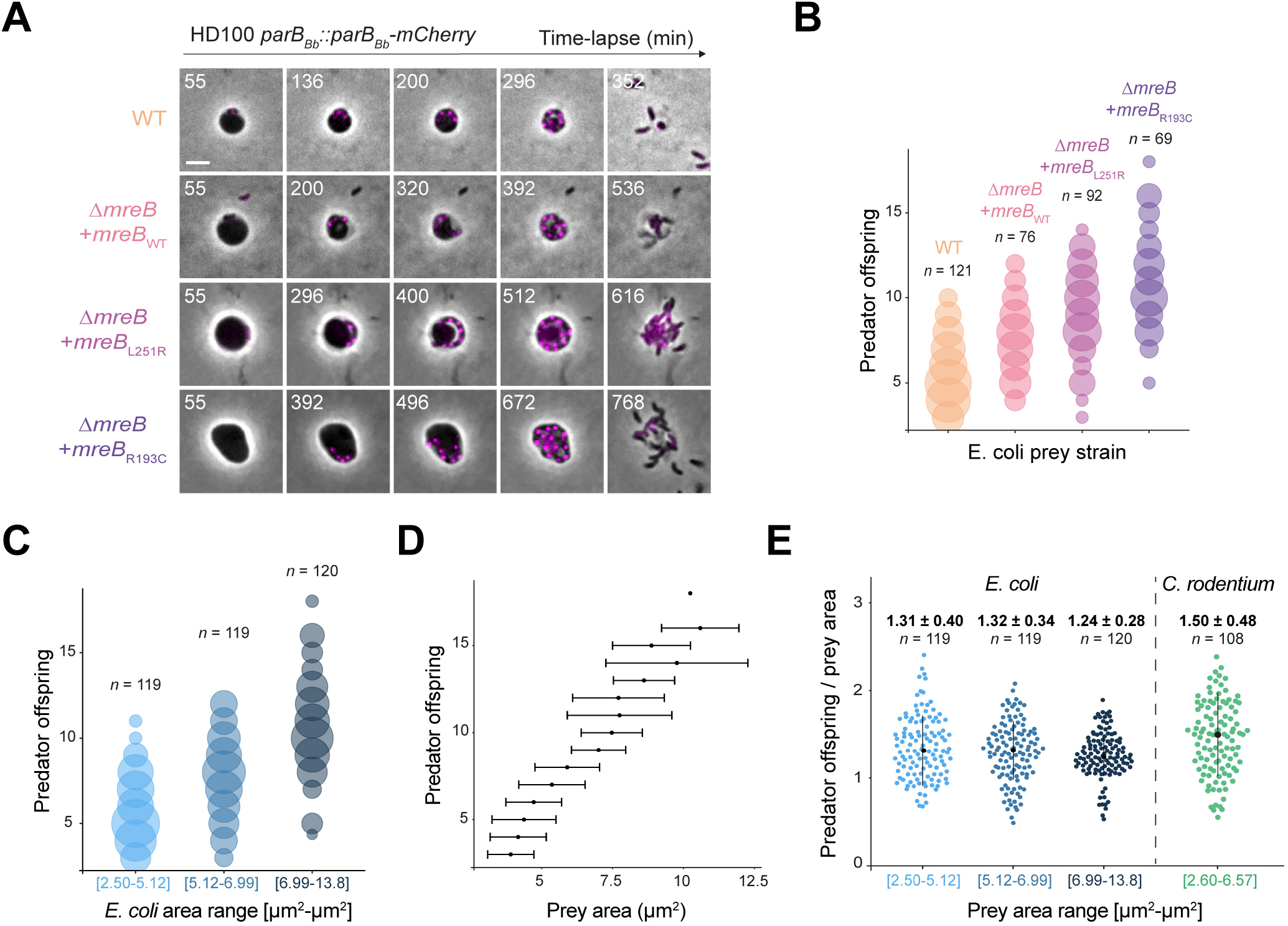
Prey size dictates the number of predator offspring. *B. bacteriovorus* carrying the chromosomal *parB*_*Bb*_*::parB*_*Bb*_*-mCherry* construct was mixed with *E. coli* MreB* prey size variants and imaged in time-lapse 55 min later at 8-min intervals. (A) ParB-mCherry foci report on the predator offspring number. Representative overlays (phase contrast and mCherry channels) of each *E. coli* variant are shown at selected timepoints, with ParB-mCherry foci displayed in magenta. Scale bar, 2 μm. (B-C) Bubble plot representation of the number of predator daughter cells (i.e., the highest number of ParB-mCherry foci per bdelloplast) recorded for each prey size variant (B) or binned by prey area in equally sized groups (C); *n*, number of analysed bdelloplasts in each group. The radius of the bubbles is proportional to the number of corresponding datapoints. (D) Distributions of single-cell prey area values corresponding to a given number of predator offspring. Median (black dot) and median absolute deviation (error bar) are shown (*n* = 358 cells including all prey sizes). (E) Beeswarm plot of the single-cell ratio between the maximum number of ParB foci and the corresponding prey area in the indicated *E. coli* size bins (left), or in *C. rodentium* (right). Values of the median (black dot), the median absolute deviation (vertical black bar) and the number of analysed bdelloplasts (*n*) are indicated above for each condition.

Providing further insight into the relationship between prey size and predator offspring numbers, the progeny number per prey area was constant over the range of tested prey cells (**Fig 1E**), with a median of 1.29 ± 0.31 μm^2^ per progeny (*n* = 358 including all prey sizes). This relationship was even conserved across prey species. For example, when we used *Citrobacter rodentium* as prey **(S3 Fig**), the progeny per prey area ratio values were similarly distributed as with *E. coli* (**Fig 1E**; median of 1.50 ± 0.48, *n* = 108). Hence, our data demonstrate quantitatively and at the single-cell level that the number of predator offspring scales with prey size through a constant relationship that is not limited to one prey species.

### Prey size modulates the duration of the predator cell cycle

We then sought to determine how prey size impacts predator offspring. Since more than one predator can simultaneously invade one prey cell [15,20], we verified that the higher progeny numbers obtained from larger preys were not simply due to more frequent co-infection of the biggest *E. coli* cells (**S1B Fig**). Such events were, in fact, rarely observed under the 1:1 predator-to-prey ratio used for all our live imaging experiments. Moreover, we noticed that the higher progeny number in larger prey cells does not result from the generation of a higher number of smaller predator daughter cells, as cell dimension distributions of newborn *B. bacteriovorus* were equal upon infection of the differently sized *E. coli* strains (**S1C Fig**). These observations imply that the final length of the filamentous mother *B. bacteriovorus* cell before division must scale with prey size.

We thus considered the possibility that the size of the infected prey determines the duration of the *B. bacteriovorus* intracellular cell cycle phase. To test this idea, we set up a tailored automated single-cell image analysis pipeline to extract quantitative insights of the predator’s growth and division within the prey (**S2 Fig**), since available image segmentation software hardly outlines the confined and entangled *B. bacteriovorus* filaments. First, we exploited the modified distribution of phase contrast pixel intensities to determine the timepoint at which the mother *B. bacteriovorus* filamentous cell visibly divides into separate daughter cells or when daughter cells start exiting the prey (two temporally close events, which for some bdelloplasts happened within the 8 min interval of our time-lapse experiment; see Methods and **Fig 2A**). We therefore consider such computed “popping time” as a relevant metric to determine when the *B. bacteriovorus* cell cycle ends, and a good proxy for when it divides. Consistent with our hypothesis, predator popping times increased with prey size, both when binned prey sizes were considered (**Fig 2B**) and at the single-cell level (**Fig 2C)**, across all *E. coli* strains. Importantly, the effect of prey size on the popping time was stronger in smaller preys than in larger preys, following a logarithmic curve (**S4 Fig**). This suggests that the *B. bacteriovorus* filament is not growing by adding a fixed volume over time, but instead elongates exponentially (see below). We thus concluded that the timing of predator cell division, i.e., the duration of its intracellular cell cycle stage, scales with the prey cell size in a non-linear way.

**Fig 2.**
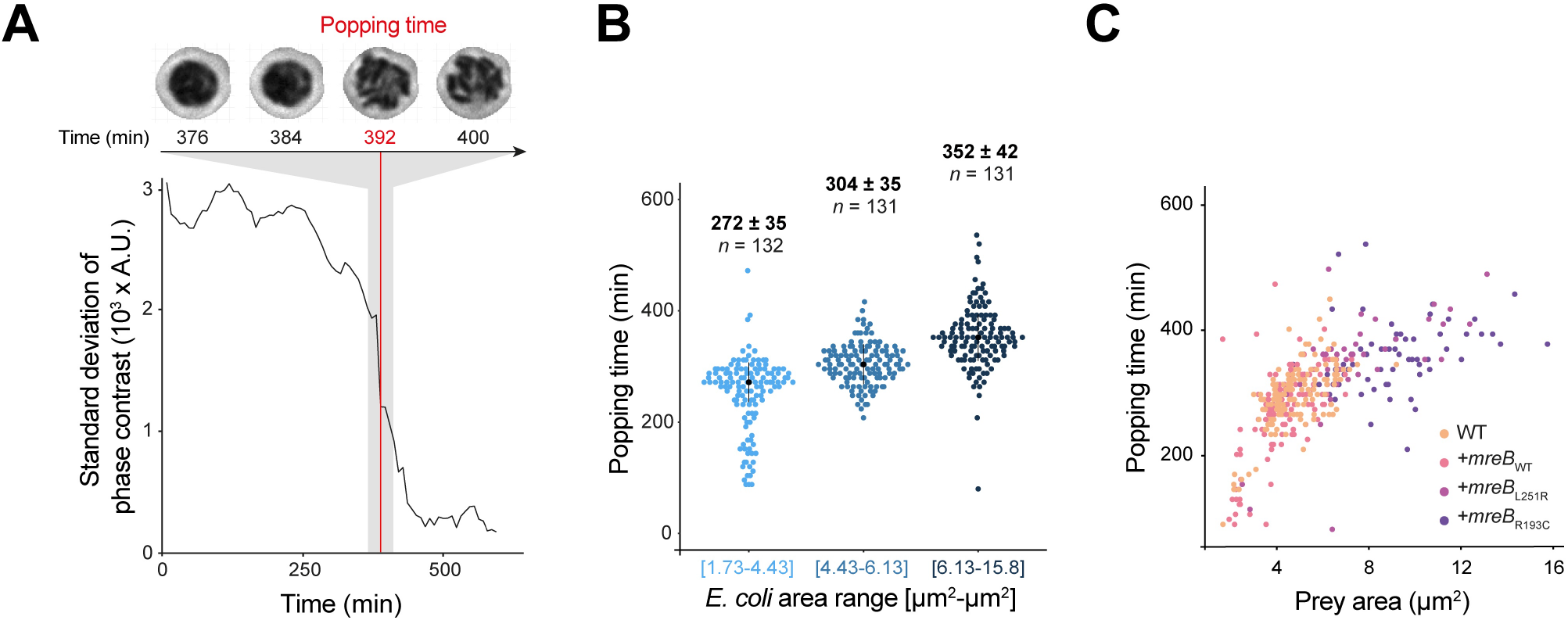
B. bacteriovorus divides later in larger prey cells. *B. bacteriovorus* (strain GL1462) was mixed with each *E. coli* MreB* prey size variant and imaged in time-lapse 55 min later at 8-min intervals. (A) Illustration of popping time determination. The standard deviation (SD) of the phase contrast is plotted over time. The timepoint corresponding to the abrupt drop of phase contrast SD is computed and the obtained popping time (vertical red line) matches with cell division of the predator filament observed by live phase contrast imaging as shown above the plot. (B) Beeswarm plot representation of the predator popping time determined for the indicated prey area categories (binned in equally sized groups). Values of the median (black dot), the median absolute deviation (vertical black bar) and the number of analysed bdelloplasts (*n*) are indicated above for each condition. (C) Scatter plot showing the relationship between popping time and prey area. Each color-coded dot represents a single predator popping time and the corresponding prey area, obtained from one *E. coli* variant (WT: *n* = 154; Δ*mreB* + *mreB*_*WT*_: *n* = 136; Δ*mreB* + *mreB*_L251R_: *n* = 41; Δ*mreB* + *mreB*_*WT*_: *n* = 63).

### Predator cells keep elongating until cell division

The observation that predator cell cycle duration is adjusted to prey size prompted us to obtain insights into the dynamics of *B. bacteriovorus* filamentous growth. To start assessing the elongation of predator cells, we took advantage of a *B. bacteriovorus* strain constitutively producing the fluorescent TdTomato protein (*Bdellovibrio*^*TdT*^) to estimate the predator cell area (a proxy for its size) based on the number of pixels above a fluorescence intensity threshold (see Methods; **S2 Fig**). This enable us to monitor the growth of single *Bdellovibrio*^*TdT*^ predator cells over time upon infection of the *E. coli* size variants by time-lapse imaging (**Fig 3A**). Single-cell growth curves confirm our interpretation that individual predator cells grow exponentially (**Fig 3B**). Moreover, the length of the *B. bacteriovorus* growth period (determined by the time at which predator elongation stops) increases with increasing prey size (**Fig 3B-D)** in a manner that is remarkably reminiscent of the relationship between cell division time and prey size (**Fig 2B and 2C**). Comparing predator popping time with growth termination time in single cells reveals the near absence of delay (median = 8 ± 12 min, *n* = 394 including all prey sizes) between the two events (**Fig 3E**), meaning that *B. bacteriovorus* cells elongate until the latest steps of cell division. In agreement with this finding, constriction (i.e., the narrowing of the cell body along the short axis that precedes cell division) becomes visible at several places along the filament while it is still growing (**S5 Fig**).

**Fig 3.**
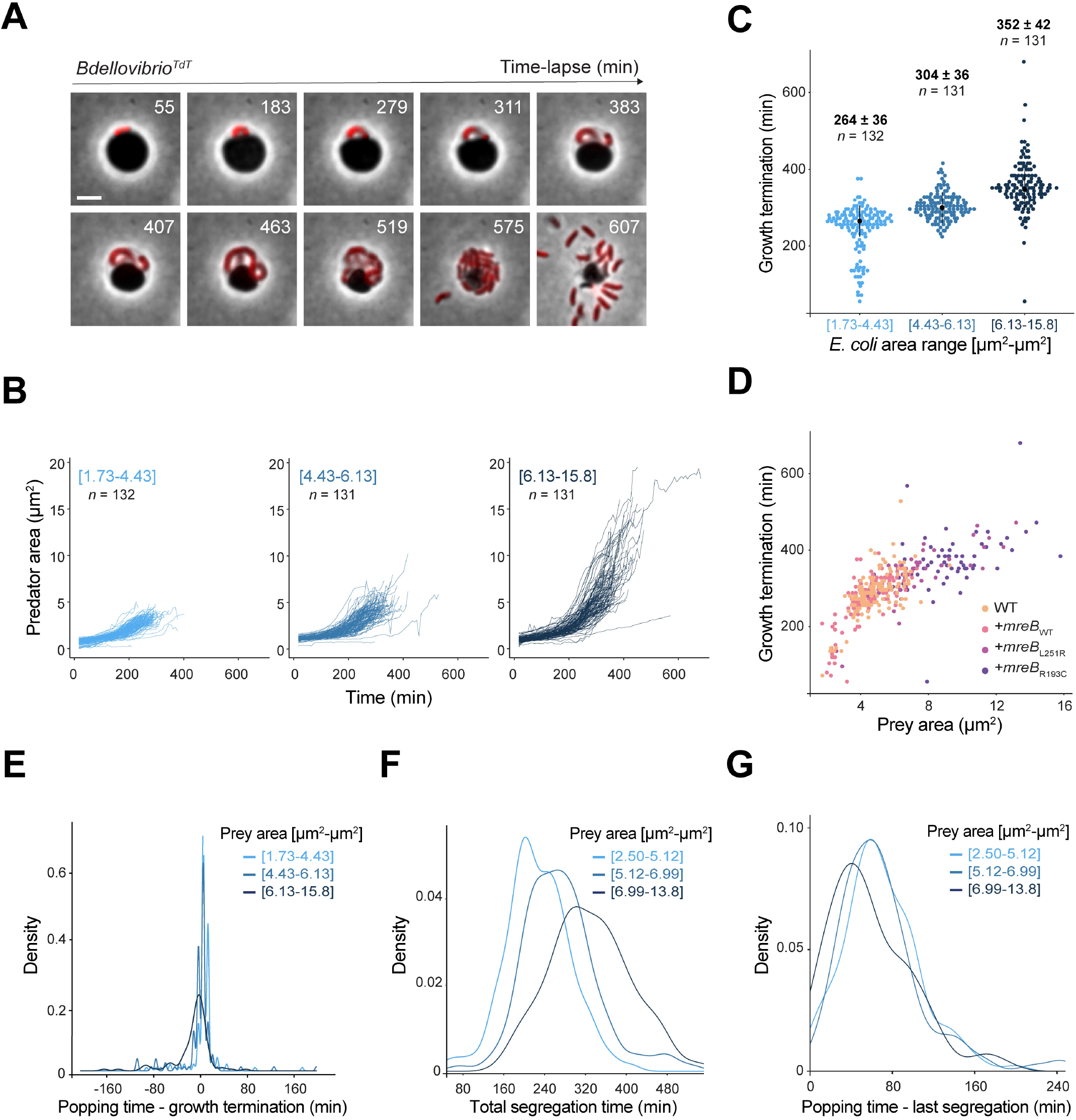
Growth of *B. bacteriovorus* relies on temporally linked cellular processes. (A-E) *B. bacteriovorus*^*TdT*^ strain was mixed with each *E. coli* MreB* prey size variant and imaged in time-lapse 55 min later at 8-min intervals. (A) TdTomato fluorescence signal is used as a reporter of *B. bacteriovorus* growth. Representative overlays (phase contrast and mCherry channels) of an infected *E. coli* cell (here MG1655 Δ*mreB* + *mreB*_*R193C*_) are shown at selected timepoints. Scale bar, 2 μm. (B) Line plots representing the growth of *B. bacteriovorus* over time (using the area of the “predator pixels” as proxy for the predator cell area, see Methods) for each binned prey area category. Each line represents the growth of one predator cell. Equally sized bins by prey area (brackets) and the number of analysed bdelloplasts (n) are indicated. (C) Beeswarm plot representation of the timepoint at which *B. bacteriovorus* stop their elongation (i.e., growth termination) determined for each prey size category (equally sized bins). Values of the median (black dot), the median absolute deviation (vertical black bar) and the number of analysed bdelloplasts (*n*) are indicated above for each bin. (D) Scatter plot showing the relationship between the growth termination time and prey size. Each color-coded dot represents the time when growth of a single *B. bacteriovorus* ends and the corresponding prey cell area, obtained from one *E. coli* variant (WT: *n* = 154; Δ*mreB* + *mreB*_*WT*_: *n* = 136; Δ*mreB* + *mreB*_L251R_: *n* = 41; Δ*mreB* + *mreB*_*R193C*_: *n* = 63). (E) Density plot of the delay between the predator growth termination time and the corresponding popping time calculated for single bdelloplasts, for each binned prey size category as indicated (brackets). The same datasets were used for B-E, with identical bins and *n* (number of analysed bdelloplasts per group) in B, C, E. (F-G) Density plots of the duration of the chromosome segregation process in *B. bacteriovorus* (carrying the chromosomal *parB*_*Bb*_*::parB*_*Bb*_*-mCherry* construct), measured in single bdelloplasts (obtained as in Fig 1) as the time between the appearance of the first ParB-mCherry focus (first segregation event) and the last ParB-mCherry focus (last segregation event) (F), or the delay between the last segregation event and the predator popping time (G) for each binned prey size category as indicated (*n* = 119 for [2.50-5.12], *n* = 119 for [5.12-6.99], *n* = 120 for [6.99-13.8]).

### The temporal coordination of predator cell cycle events is robust across prey sizes

Strikingly, the temporal proximity between the end of predator cell elongation and division is unaffected by prey size (**Fig 3E)**, hinting that the link between these key processes is wired in the cell cycle. Because cell cycle progression is guided mainly by chromosome replication and segregation in other species [21,22], we wondered how *B. bacteriovorus* (which combines multiple asynchronous rounds of replication with progressive segregation [17]) integrates these complex processes within cell cycles of various durations. To grasp predator chromosome dynamics in differently sized preys, we extracted the duration of the entire chromosome segregation process by recording when the first and the last segregation events occur (corresponding to the appearance of the first and last ParB focus, respectively). In line with the predator cell cycle duration adjustment, the segregation period scaled with prey size (**Fig 3F)**. In addition, the delay between the last segregation event and the predator popping time (median = 64 ± 36 min, *n* = 358 including all prey sizes) was not visibly affected by prey size (**Fig 3G**), suggesting that the time required to complete division upon the proper segregation of sister chromosomes is fixed. Conveniently, the segregation period also reflects the chromosome synthesis period (called C period) since (i) the first ParB focus follows the initiation of the first chromosome replication round in *B. bacteriovorus* (by 41 min on average [17]) and (ii) no new chromosome replication round starts after the last ParB focus is detected [17]. Altogether, our data indicate that chromosome dynamics, cell growth, and cell division are coupled in a manner that appears wired in the cell cycle: while the total duration of the proliferative phase until division (including the chromosome-related processes and cell growth) adjusts to prey size, the time interval between these processes is invariable.

### The nutritional quality, but not the size of the prey cell, modulates the predator growth rate

Until now, we have modified the size of the prey, which altered both the total resources and the space available to accomodate *B. bacteriovorus* filamentation. To refine our understanding of the impact of the prey on predator physiology, we set out to modify the composition of the prey cell and test its impact on the proliferation of predator cells. To this end, we grew *E. coli* in different media, either LB (a rich and undefined medium, as done in **Fig 1-3**) or M9 supplemented with glucose and casamino acids (a relatively poor and defined medium) prior to infection by *B. bacteriovorus*. Indeed, previous work demonstrated that in these distinct growth conditions, the cellular dry mass of *E. coli* does not change [23,24] but the cellular resource allocation (i.e., the macromolecular composition of the cytoplasm) varies [25–27]. For instance, *E. coli* cells grown in LB contain relatively more proteins than cells grown in M9. Strikingly, modifying the prey cell composition did not considerably impact the output of the proliferation stage, i.e., the number of *B. bacteriovorus* daughter cells produced per “unit” of prey (**Fig 4A**). However, the ratio between predator popping time and prey area, a metric of predator cell cycle duration per prey size unit, was higher in M9-than in LB-grown prey (**Fig 4B**). This indicates that, at comparable prey size, the time required to achieve a given progeny number was longer in M9-grown prey.

**Fig 4.**
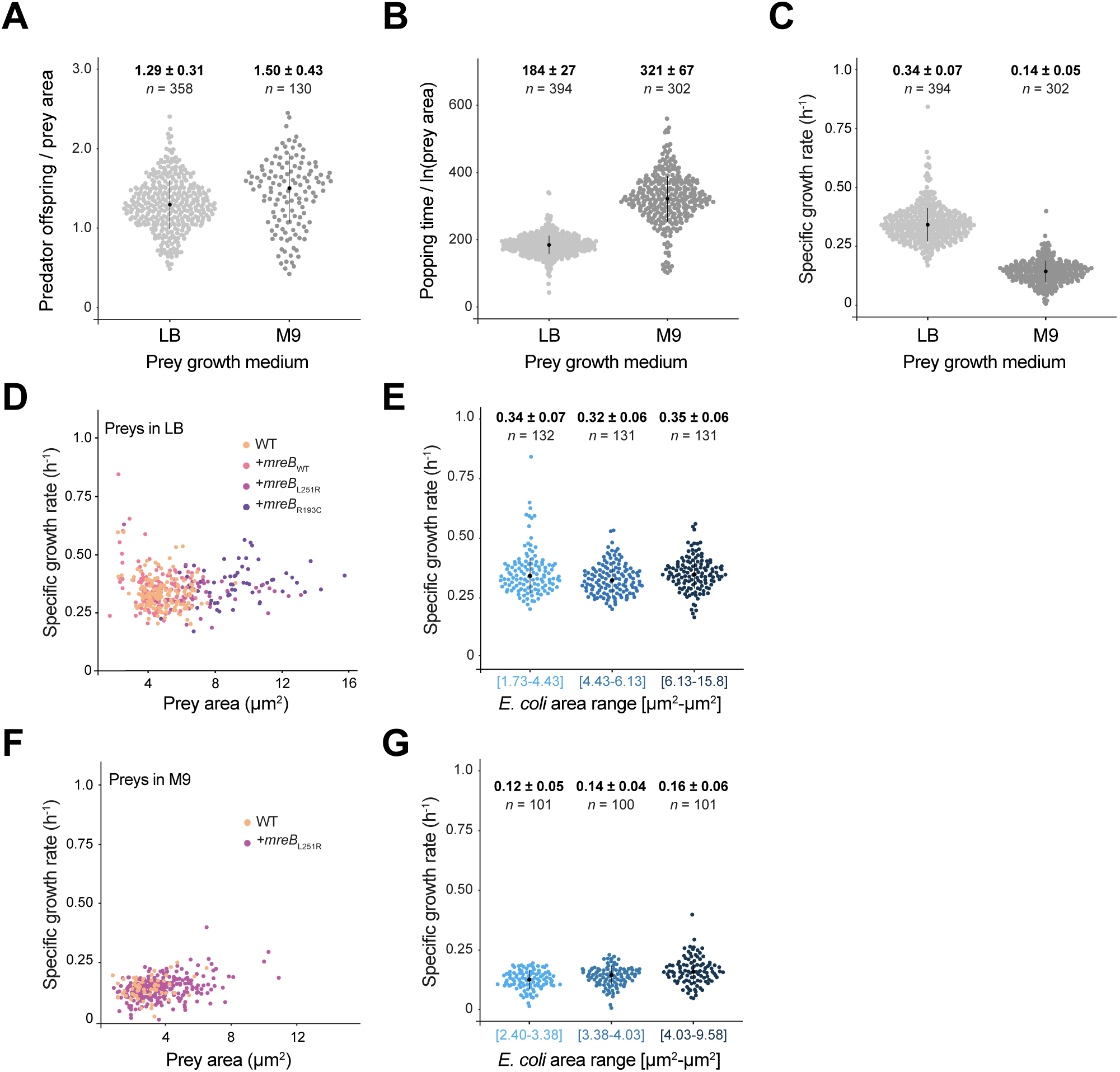
The nutritive quality of the prey influences the specific growth rate of predators. *B. bacteriovorus* carrying the chromosomal *parB*_*Bb*_*::parB*_*Bb*_*-mCherry* construct (A) or producing cytoplasmic TdTomato (B-G) was mixed with each *E. coli* MreB* prey size variant (grown in LB), or the WT or Δ*mreB* + *mreB*_L251_ (grown in M9) and imaged in time-lapse 55 min later at 8-min intervals. (A-B) Beeswarm plot of the single-cell ratio between the maximum number of ParB foci and the corresponding prey area (A) or the single-cell ratio between popping time and the corresponding logarithm of the prey area (B), when prey cells were grown in LB or in M9 medium prior infection as indicated, including all prey sizes. In (B), the ln(area) was used since predator popping time and prey area values display a logarithmic relationship (Fig 2C and S4 Fig). Values of the median (black dot), the median absolute deviation (vertical black bar) and the number of analysed bdelloplasts (*n*) are indicated for each condition. (C) Beeswarm plot representation of the *B. bacteriovorus* specific growth rate (sGR) determined during infection of *E. coli* size variants previously grown in either LB or M9 medium. Values of the median (black dot), the median absolute deviation (vertical black bar) and the number of analysed bdelloplasts (*n*) are indicated above for each condition. (D) Scatter plot showing the single-cell relationship between the *B. bacteriovorus* specific growth rate (sGR) and prey size. *E. coli* cells size variants were grown in LB prior infection. Each color-coded dot represents the sGR of a single *B. bacteriovorus* cell and the corresponding prey area, obtained from one *E. coli* strain (WT: *n* = 155; Δ*mreB* + *mreB*_*WT*_: *n* = 136; Δ*mreB* + *mreB*_L251R_: *n* = 41; Δ*mreB* + *mreB*_*WT*_: *n* = 63). (E) Beeswarm plot representation of the *B. bacteriovorus* sGR determined for the indicated prey size categories (equally sized bins, same cells as in Fig 4D). Values of the median (black dot), the median absolute deviation (vertical black bar) and the number of analysed bdelloplasts (*n*) are indicated for each condition. (F) Scatter plot showing the relationship between the *B. bacteriovorus* specific growth rate (sGR) and prey size. WT and the Δ*mreB* + *mreB*_L251R_ *E. coli* strains (which together cover a relatively large range of prey size) were grown in supplemented M9 prior synchronized predation by *B. bacteriovorus*. Each color-coded dot represents the sGR of a single *B. bacteriovorus* cell and the corresponding prey area, obtained from one *E. coli* strain (WT: *n* = 58; Δ*mreB* + *mreB*_L251R_: *n* = 244). (G) Beeswarm plot representation of the *B. bacteriovorus* sGR determined for the indicated prey size categories (equally sized bins, same cells as in Fig 4F). Values of the median (black dot), the median absolute deviation (vertical black bar) and the number of analysed bdelloplasts (*n*) are indicated above for each condition. (F-G) Beeswarm plot of the single-cell ratio between the maximum number of ParB foci and the corresponding prey area (F) or the single-cell ratio between popping point and the corresponding logarithm of the prey area (G), when prey cells were previously grown in LB or in M9 medium, including all prey sizes. In (G), the ln(area) was used since predator popping point and prey area values display a logarithmic relationship (Fig 2C and S4 Fig). Values of the median (black dot), the median absolute deviation (vertical black bar) and the number of analysed bdelloplasts (*n*) are indicated for each condition.

Consistent with these findings, predator cells elongate slower in M9-grown prey than in LB-grown prey, as the specific growth rate of *B. bacteriovorus* (computed from single-cell growth curves obtained as described above, see Methods) was about two-fold lower for predators feeding on M9-grown *E. coli* (median: 0.14 ± 0.05.h^-1^, *n* = 302) compared to predators in LB-grown prey (0.34 ± 0.07.h^-1^, *n* = 394) (**Fig 4C**). Thus, the cellular content of the prey, likely through its macromolecular composition, is a key factor that constrains the intracellular proliferation of predatory bacteria. In bacteria, the growth rate depends on the nutritional quality of the medium: bacteria grow faster in “rich” medium (containing higher concentration of metabolizable nutrients) than in “poor” medium [28–31]. In this case, the composition of LB-or M9-grown prey can be considered as a “rich” or “poor” medium for *B. bacteriovorus*, respectively, even though these cells contain the same total amount of resources. Accordingly, the cell-to-cell variability in predator growth rate was independent of prey size, regardless of the nutritional quality of the prey (**Fig 4D-G**). This suggests that the size of the prey cell, and therefore the total resources and/or the space within the prey, do not impact the speed of growth, while they impact the duration of the cell cycle (**Fig 2 and 3**) under the tested conditions.

Altogether, our data hint that the prey nutritive quality steers the speed of predator growth, while the size of the prey cell eventually determines the number of predator offspring.

## Discussion

In this study, we examined how the size and composition of the prey modulates the proliferation of endobiotic predators. By monitoring key aspects of the *B. bacteriovorus* growth phase in prey cells of different size and content, we provide important insights into the temporal control of the intracellular phase of the predator cell cycle.

First, we present quantitative evidence for the effect of prey size on the *B. bacteriovorus* lifecycle. The idea that the number of predator daughter cells is somehow related to the size of the prey had been proposed previously but lacked unambiguous and single-cell demonstration [15,18]. Besides, the impact of prey size on *B. bacteriovorus* cell cycle progression had never been explored. We have tackled those questions systematically, by taking advantage of (i) characterized mutants within the same *E. coli* genetic background [19], allowing reliable interpretation of size-dependent effects; (ii) the automated monitoring of an established marker of the *B. bacteriovorus* chromosomal *ori* [17], which reports on the timing of chromosome-related events but also on the number of predator progeny; (iii) a cytoplasmic fluorescence signal as a proxy to estimate the area of individual predator cells; and (iv) a tailored image analysis pipeline that extracts prey size and predator features at the single-cell level from time-lapse images. Our results not only demonstrate that the number of *B. bacteriovorus* offspring is determined by the size of the prey cell, but they also show that the relationship between prey area and progeny number in single cells is conserved across prey species and prey nutritive quality. This finding strongly supports the idea that the *B. bacteriovorus* cell cycle is wired to maximize usage of the prey resources and space.

Secondly, we reveal that modifying the content or “resource allocation” of the prey cell (by changing their growth medium), modulates the growth rate of predators. This is remarkably reminiscent of the impact of nutrients richness on the growth rate of model species. Furthermore, the predator growth rate was constant across prey sizes, highlighting that the nutritional quality, but not the total resources available within the prey, determines the speed of *B. bacteriovorus* elongation. Our findings also provide clues on what prey molecules *B. bacteriovorus* preferentially feeds upon. Because *E. coli* cells growing in rich medium (e.g., LB) contain relatively more (ribosomal) RNA and proteins than in poor medium (e.g., M9) [25–27,32], RNAs and/or proteins might constitute the primary resources fueling predator growth. With the impressive number (∼150) of putative proteases/peptidases encoded in its genome [33], *B. bacteriovorus* seems particularly well adapted to this diet.

Based on our results, we propose a model in which the duration of the intracellular proliferative phase of the *B. bacteriovorus* cell cycle depends on two factors (**Fig 5**): (i) the size of the prey, which determines how many daughter cells are produced, and therefore how much the predator cell needs to grow and how many copies of its chromosome must be synthesized and segregated **(Fig 5A**), and (ii) the nutritive quality of the prey cell, which impacts how fast the predator elongates (**Fig 5B**). The resulting outstanding question is how and when the mother predator “decides” to stop its growth phase and initiate division. Presumably, such a drastic developmental switch must be finely regulated in time: dividing too early might imply wasting available resources, dividing too late could be associated with the abortion of ongoing cellular processes, which could be lethal for the progeny. As previously proposed [15,34,35], we envision that diffusible prey-derived molecules, such as secondary metabolites, report on the available resources and serve as a not-to-be-missed “stop signal” for *B. bacteriovorus*. Alternatively, or in addition, mechanical constraints on the growing filament (due to space reduction within the bdelloplast) might also constitute a signal to end the predator proliferative phase. The nature of the prey-derived factor(s) is still mysterious, but our results indicate that the “stop signal” is integrated upstream the chain reaction of key events that take place in the predator cell (i.e., finalize DNA synthesis and segregation, halt cell growth and complete cell division; **Fig 5A**). We think that temporally coupling these late steps of the proliferative phase is advantageous for *B. bacteriovorus*: only one process would need to react to the “stop signal” and all downstream events would be launched one after the other with an invariable timing – like a domino effect. Another appealing implication of our findings is that the “stop signal” likely occurs before the complete depletion of nutrients within the prey, as it must be perceived well before the arrest of growth and the completion of cell division.

**Fig 5.**
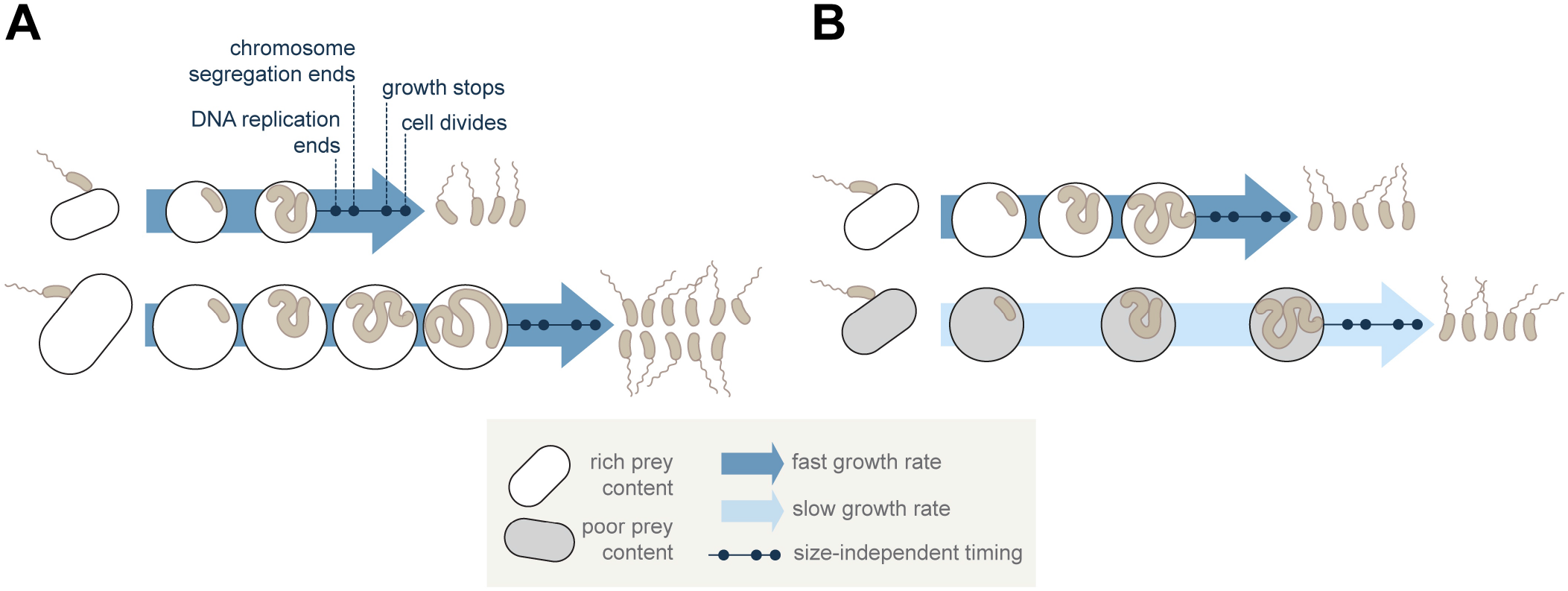
Impact of prey size and content on the intracellular cell cycle phase of *B. bacteriovorus* predators. (A) *B. bacteriovorus* adjusts the duration (but not the speed) of its growth phase to the size of the prey. As a result, the number of predator progeny scales with prey size. Time intervals between cell cycle events (e.g. end of growth to cell division, end of chromosome replication and segregation cycles to cell division) are not affected by prey size variations. (B) The specific growth rate of *B. bacteriovorus* depends on the prey nutritive quality: at the same cell size, prey cells grown in rich medium will accommodate faster growing predator cells than prey cells grown in poor medium.

In model species, cell cycle progression is mainly controlled by chromosome-related processes, under normal conditions [21,22] and upon stress (e.g., to licence the stepwise release of “repaired” daughter cells from filamentous *E. coli* recovering from DNA damage [36]. In this respect, we can consider the depletion of prey resources and space as a stress encountered by *B. bacteriovorus* – here as part of its normal predatory cell cycle. Along those ideas and based on our results and previous work [17], we propose that fundamental chromosome-related process such as replication and segregation are major checkpoints regulating the length of the predatory cell cycle. In this scenario, the “stop signal” might prevent the initiation of new rounds of DNA replication, thereby controlling the duration of the C period in *B. bacteriovorus* and, in turn, the total duration of chromosome segregation and the time when cell division completes. The initiation of cell constriction while the *B*. bacteriovorus is still elongating likely contributes to optimize the predator intracellular residency time. Our data also hint that cells keep growing after the final number of progeny is set (i.e., when the last ParB focus appears), perhaps to adjust the size of future siblings. Identifying how *B. bacteriovorus* regulates the activity and positioning of the growth and division machineries is an exciting prospect that should reveal additional insights into the complex spatiotemporal coordination of the predatory cell cycle.

From a general point of view, one may ask whether filamentous growth and synchronous division, as opposed to successive binary reproduction, is a favourable strategy to thrive in a confined micro-environment. We speculate that the peculiar proliferation adopted by *B. bacteriovorus* comes with the benefit of centralizing sensing and responses to prey cues in one single cell, facilitating the cell cycle modulation that we have observed. Besides, successive binary divisions would likely impede predator proliferation, as the resulting heterogenous progeny (e.g., growing cells vs newborn predators ready to escape) might threaten the integrity of the closed nest. We therefore envision that endobiotic predators evolved their mode of growth and division to enhance their capacity to exploit prey cells from the inside. Our data strongly suggest that the intriguing proliferation of *B. bacteriovorus* is built on the same general principles observed in binary-dividing bacteria, e.g., adjusting growth rate to nutrient quality and tightly coordinating cell cycle events. Only here, we find the remarkable twist of fine-tuning the length of the proliferative phase of the cell cycle to the size of the nest in which it takes place. Thus, the *B. bacteriovorus* cell cycle features both robustness and adaptability to the variability of its prey. Finally, our study illustrates that studying non-canonical lifestyles has the potential to reveal elaborate strategies employed by bacteria to prosper in all possible settings.

## Material and methods

### Bacterial strains, growth conditions and chemicals

All strains used in this study are listed in Table S1. *Bdellovibrio bacteriovorus* HD100 (taxon: 264462) was used as a model predator strain. *Escherichia coli* MG1655 derivatives and *Citrobacter rodentium* (taxon: 67825) were used as prey. *E. coli* and *C. rodentium* cells were routinely grown in Lysogeny Broth (LB, MP Biomedicals) or, when indicated, in M9 minimal medium (mpbio^™^) supplemented with glucose 0.4%, casaminoacids 0.2%, vitamin B1 1 μg.mL^−1^, MgS0_4_ 1 mM and CaCl_2_ 0.3 mM at 37°C with aeration. Plasmids were maintained by adding kanamycin (50 μg.mL^−1^) or chloramphenicol (15 μg.mL^−1^). *B. bacteriovorus* strains were routinely grown in DNB medium (Dilute Nutrient Broth, Becton, Dickinson and Company, supplemented with 2 mM CaCl_2_ and 3 mM MgCl_2_ salts) with *E. coli* MG1655 as prey at 30°C and constant shaking as previously described [37].

**Table S1.**
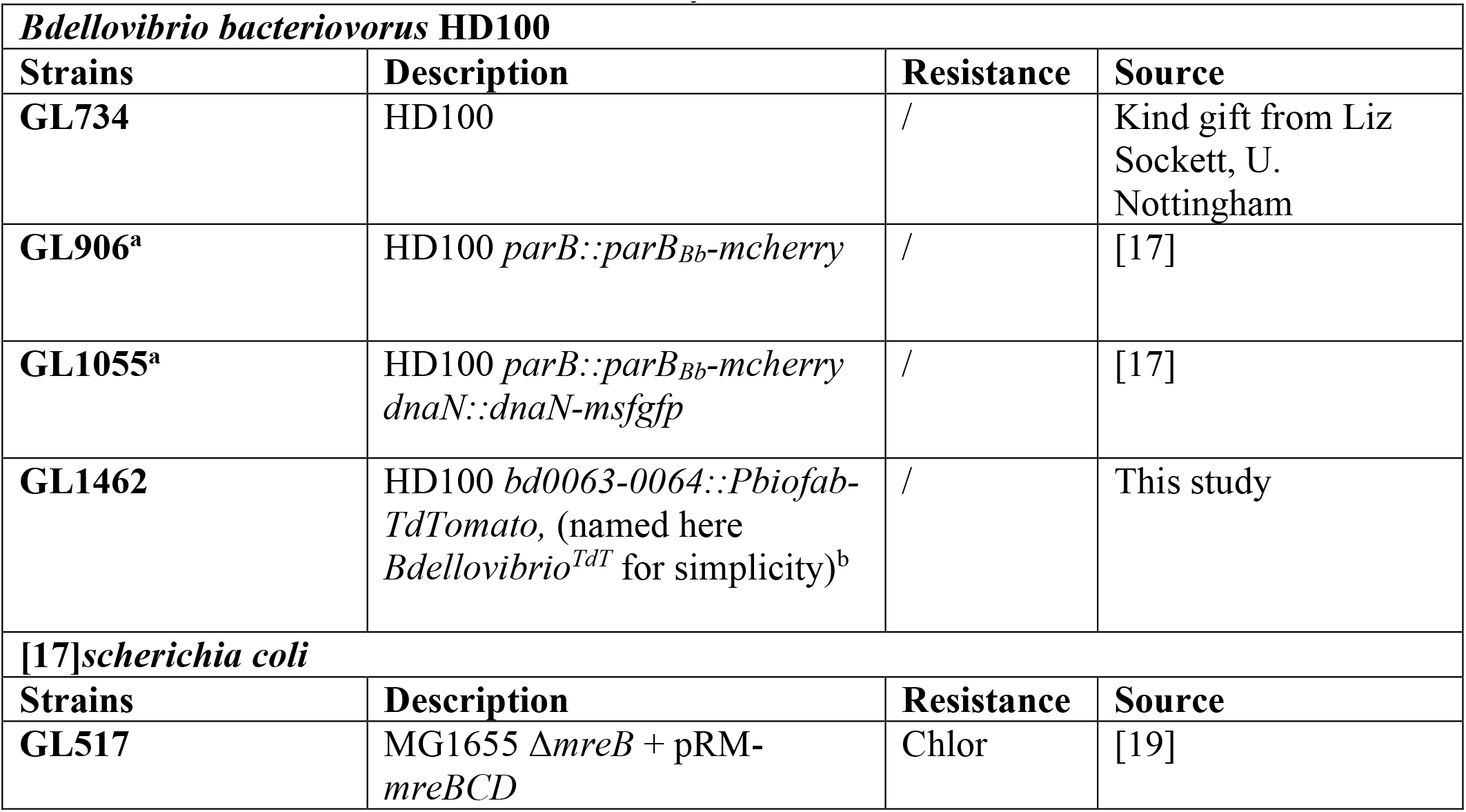
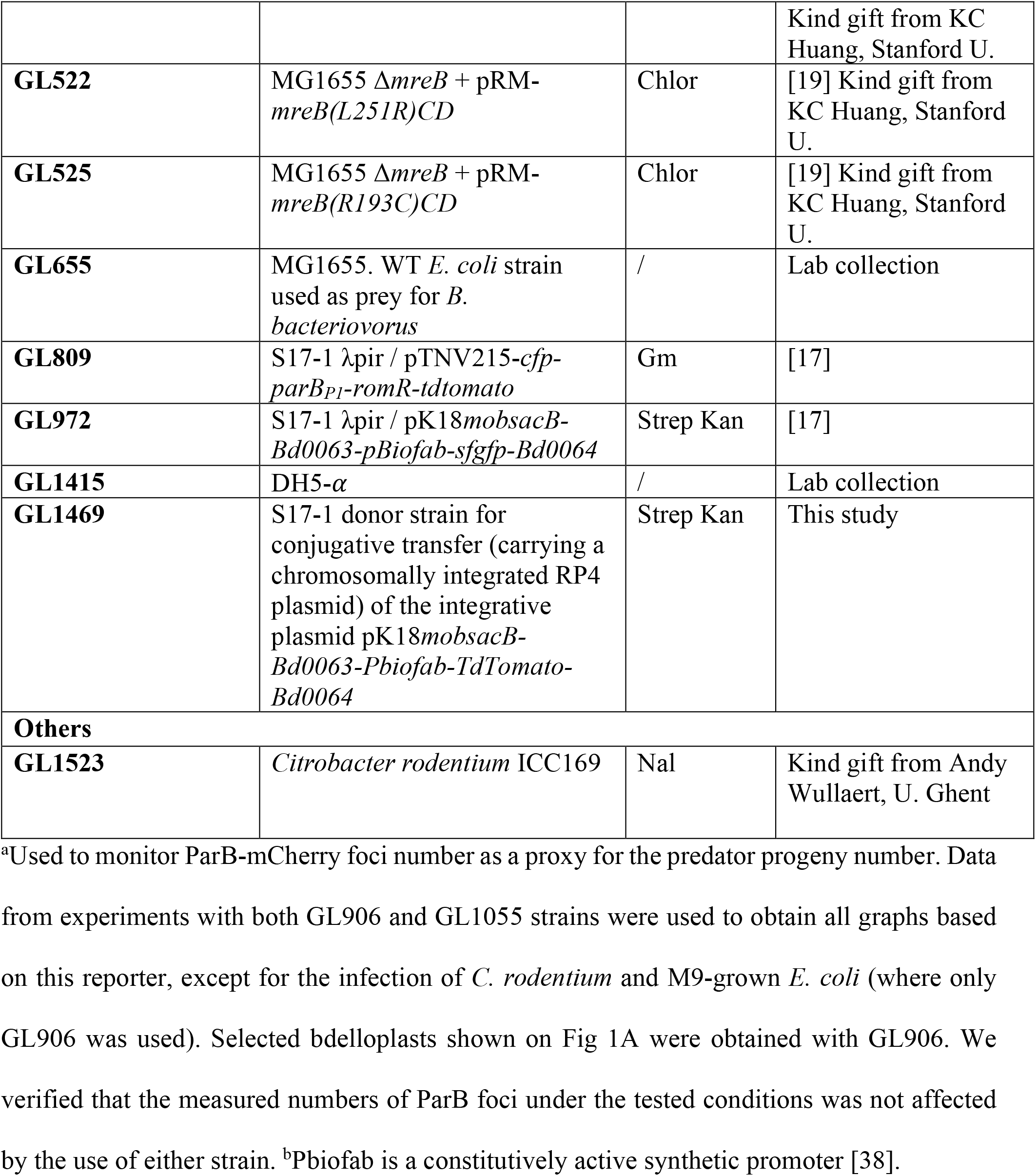
Bacterial strains used in this study.

### B. bacteriovorus strain construction

All primers used in this study are listed in Table S2. Standard molecular cloning methods were used, and DNA assembly was performed using the NEBuilder HiFi mix (New England Biolabs). *B. bacteriovorus bd0063-0064::Pbiofab-TdTomato* (named *Bdellovibrio*^*TdT*^ for simplicity) was generated from the wild-type strain HD100 by mating between a *E. coli* S17-1 donor strain (GL1459) carrying the mobilizable plasmid pK18mobsacB-Bd0063-Pbiofab-TdTomato-Bd0064 (see construction method below) and *B. bacteriovorus* HD100 as the recipient strain. Briefly, exponentially growing-donor cells were harvested and washed twice in a DNB-salts medium before resuspension in 1:10 of the initial volume in DNB-salts. This donor suspension was mixed at equal volume with a fresh overnight lysate of the recipient strain. The mating mix was incubated for a minimum of 24 h at 30°C with shaking before plating on a selective medium using the double-layer technique (ref). Single plaques were isolated and transconjugants were confirmed by microscopy, PCR, and sequencing. Scarless allelic replacement into the HD100 chromosome was performed using a strategy based on the two-step recombination with a pK18mobsacB-derived suicide vector as in [39], screened by PCR and verified by DNA sequencing. To construct the mobilizable and integrative plasmid pK18mobsacB-Bd0063-Pbiofab-TdTomato-Bd0064, the pK18mobsacB-Bd0063Pbiofab-sfgfp-Bd0064 plasmid (in GL972) was PCR amplified with primers oGL967/oGL783 to remove the *sfgfp*-coding sequence. The *tdtomato*-coding sequence was then amplified with oGL887/oGL968 from the pTNV215-*cfp-parBP1-romR-tdtomato* plasmid (in GL809) and cloned in the opened vector by DNA assembly.

**Table S2.**
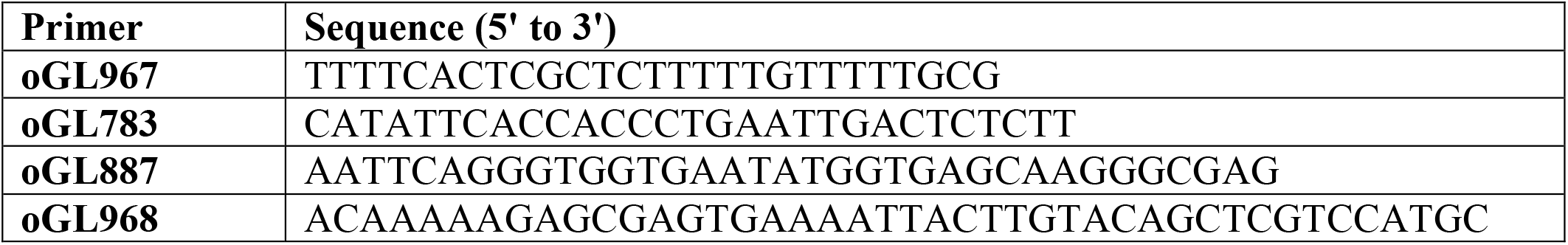
Primer information.

### Live-cell imaging

*B. bacteriovorus* strains were first grown overnight on WT MG1655 (as described [37]) before the start of the imaging experiment. For time-lapse imaging of synchronous predation cycles, *E. coli* prey cells were grown in LB medium (or supplemented M9 when indicated) to late exponential phase (OD_600_ = 0.8), harvested at 2600 x *g* at room temperature (RT) for 5 minutes, washed twice and resuspended in DNB-salts medium. Then, *B. bacteriovorus* and *E. coli* were mixed at a 1:1 volume ratio to allow most prey cells to be infected simultaneously or with a 1:1, 2:1, and 5:1 ratio for a co-infection assay when indicated. We consider the prey-predator mixing step as the time 0 in all our synchronous predation imaging experiments. Cells were co-incubated for 45 min to 1 h at 30°C prior to imaging on 1.2% DNB-agarose pads. In time-lapse experiments, the same fields of view on the pad were imaged at regular intervals as indicated, with the enclosure temperature set to 28°C. For snapshots of fresh (newborn) attack phase *B. bacteriovorus* (**S1C Fig**), cells were spotted on 1.2% agarose pads prepared with DNB-salts medium after one full synchronous predation cycle performed as described above. For snapshots of *E. coli* strains (**S1A Fig**), overnight cultures were diluted at least 1:200, grown to exponential phase, and washed twice in DNB-salts before spotting on 1.2% agarose pads prepared with DNB-salts medium.

### Image acquisition

Phase contrast and fluorescence images were acquired on a fully motorized Ti2-E inverted epifluorescence microscope (Nikon) equipped with a CFI Plan Apochromat *λ* DM 100x 1.45/0.13 mm Ph3 oil objective (Nikon), a Sola SEII FISH illuminator (Lumencor), a Prime95B camera (Photometrics) or a PrimeBSI camera (Photometrics), a temperature-controlled light-protected enclosure (Okolab), and a mCherry-C filter cube (32 mm, excitation 562/40, dichroic 593, emission 640/75; Nikon) to image TdTomato and ParB-mCherry. Multi-dimensional image acquisition was controlled by the NIS-Ar software (Nikon). Pixel size was 0.074 μm for the Prime95B camera (using the 1.5X built-in zoom lens of the Ti2-E microscope) or 0.065 μm for the PrimeBSI camera (with 1X zoom lens). Identical LED illumination power and exposure times were applied when imaging several strains and/or conditions in one experiment and across replicates of the same experiment. They were set to the minimum for time-lapse acquisitions to limit phototoxicity.

### Image processing

For figure preparation, images were processed with FIJI [40] keeping contrast and brightness settings identical for all regions of interest in each figure. Figures were assembled and annotated using Adobe Illustrator (Adobe Inc.).

### Co-infection quantification

Co-infections were defined as events in which more than one *B. bacteriovorus*^*TdT*^ predator is present inside an *E. coli* prey cell upon 1 h post-infection. Live-cell imaging was performed as mentioned above. To avoid counting predators attached or close to the prey envelope, time-lapse recordings were performed, and only predators for which growth was observed (i.e., predators feeding inside their prey) were considered (**S1B Fig**).

### Bdelloplast detection from phase contrast images

Images were aligned on the first phase frame using the Image Stabilizer ImageJ plugin [41]. As the position and shape of the rounded infected prey remained unchanged during the time-lapse experiments, only the outlines of the first frame (i.e., the first timepoint) of each movie were detected and subsequently reused for all timepoints (**S2 Fig**). Subpixel detection of the bdelloplasts was done with the cellDetection tool in the open-source image segmentation software Oufti [42]. To avoid the misdetection of ParB foci and to compute the popping time (see below), the bdelloplast outlines were expanded by a buffer region of 0.5 μm.

### Prey size measurement

As the *B. bacteriovorus* invasion step is not recorded in our time-lapses, the size of each infected *E. coli* cell was approximated by the measured area (in μm^2^) of the corresponding bdelloplast outline obtained at subpixel resolution from the first frame as described above (**S2 Fig**). In addition, the area of uninfected *E. coli* (**S1A Fig**) was also computed from subpixel outlines obtained with Oufti [42]. When indicated, bdelloplasts were split into 3 bins containing the same number of cells along the distribution of area values. We chose to use cell area as a proxy for assessing cell size as this is the most precise measurement, we could extract from 2D images without making assumptions about the third dimension and the actual shape of the cells. While this is a potential limitation of our study inherent to the imaging setup, volume approximations would have decreased the precision of our method. Of note, estimations of ellipsoid volumes (considering that cells are immobilized between the agarose pad and the coverslip) of various heights and based on the measured range of bdelloplast radius indicate little difference in absolute values of such approximated volumes and the corresponding bdelloplast areas, supporting the relevance of our approach.

### Foci detection from fluorescence images

ParB-mCherry foci were detected from the fluorescence images using the LoG detector from the TrackMate ImageJ plugin [43] using the following set of parameters: estimated object diameter = 3.5 pixels; quality threshold = 15; with sub-pixel localization. Detected foci were then further filtered based on their contrast and signal-to-noise ratio. To properly define the appearance of the first ParB-mCherry focus, bdelloplasts infected by predators exhibiting foci before the fourth frame was not considered. The beginning of the first or last chromosome segregation event was defined as the first frame in the time-lapse (i.e., the first timepoint) where the first ParB-mCherry focus or the maximum number of ParB-mCherry foci is detected, respectively (**S2 Fig**).

### Popping time determination

The popping time corresponds to the timepoint at which a *B. bacteriovorus* filament divides into daughter cells (**S2 Fig**). In phase contrast time-lapses, there is a sharp decrease in the contrast between the bdelloplast and its surroundings when *B. bacteriovorus* divides (**Fig 2A**). This was used as a proxy to determine the popping time. Next, the contrast was computed as the weighted standard deviation of the phase contrast pixels within the buffered bdelloplast outline. Here each pixel was weighed by its coverage fraction within the bdelloplast outline. Finally, the maximum decrease, in contrast, was computed as the maximum difference of two successive running means of the contrast values. We verified manually that the popping time computed by this method corresponded to the visible division of the mother predator cell on a subset of bdelloplasts across all prey strains. When the temporal resolution of our time-lapse experiment (8 minutes) did not allow us to catch the division event before daughter cells start exiting the prey (two events following each other closely in time, based on our observations and [15,16]), the popping time then corresponded to an already initiated release of newborn predators.

### Growth curve extraction and growth rate calculation

The fluorescence signal from the constitutively produced TdTomato in the *Bdellovibrio*^*TdT*^ strain was used as a proxy to quantify predator elongation over time (**S2 Fig**). To separate true signal from background noise, a threshold was set in each bdelloplast as

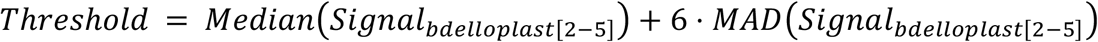

where *MAD* stands for median absolute deviation, *Signal*_*bdelloplast[2-5]*_ is the array of fluorescence intensity values of all pixels inside the given buffered bdelloplast outline in frames 2 to 5. The predator area was subsequently computed as the number of pixels above the threshold value converted to μm^2^. The initial predator cell area obtained by this method falls within a range that matches previous measurements based on automated segmentation of attack-phase *B. bacteriovorus* cells [4]. Overlays of the fluorescence channel image with the selected pixels indicate a satisfying approximation of the position and area of the *B. bacteriovorus* cell across all *E. coli* size variants and throughout the time-lapse experiments (**Supplementary Movie 1**), validating our approach.

To compute the specific growth rate (*a*) of each *B. bacteriovorus* cell, a linear model (*y*=*ax*+*b*) was fitted on the log transformation of the area (*y*) over the time (*x*). Timepoints after the end of predator growth in each bdelloplast were removed, where the end of growth was defined as the first local maximum in the calculated *B. bacteriovorus* area over time. A fraction of the detected bdelloplasts contained predator cells that did not grow or did not finish their growth cycle before the end of the time-lapse movie. Three filters were applied to discard these bdelloplasts from the analysis when any of the following conditions was met: (i) detection of pixels above the defined threshold for less than 5% of the whole time-lapse; (ii) mean predator filament size (calculated from the predator pixel area at all timepoints) lower than 1.5 times their initial size; (iii) *B. bacteriovorus* filaments with a final area lower than 1.95 times their mean area. Initial predator sizes are evenly distributed across prey size bins (**Fig. 3B**), indicating the absence of prey-dependent bias in our estimation of the predator pixel area.

### Pipeline analysis and data availability

The image analysis pipeline used in this study is described schematically in **S2 Fig**. Foci detection, growth curve extraction, and popping times detection were combined in a configurable Rmarkdown file available on https://github.com/geraldinelaloux/prey_size_analysis with the required documentation for installation and running. Parameters used to obtain the plotted https://github.com/geraldinelaloux/prey_size_analysis/Experiments.

### Statistical analyses

All analyses of microscopy images were performed using several representative fields of view from at least three independent biological replicates. Means and standard deviations (SD), medians and median absolute deviations (MAD), and linear models were calculated in R using the base R functions. Pearson correlation was calculated in R using the corrr package [44]. Density plots in Fig 3E-G were computed as kernel density estimates and plotted with the ggplot2 function geom_density [45].

## Acknowledgements

We are grateful to Charles de Pierpont for constructing the GL1462 *Bdellovibrio* strain and for excellent technical support, all members of the Laloux lab and Michaël Deghelt for stimulating discussions and critical review of the manuscript, Sander Govers for advice on bacterial growth and for critical reading of the draft, K.C. Huang for kindly providing the *mreB\** mutants, Andy Wullaert for sharing the *Citrobacter rodentium* strain, the team of Eric Cascales for insightful discussion, and Karl Kochanowski for advices about bacterial growth metrics.

This work was supported by the European Commission (ERC Starting Grant PREDATOR #802331). Y.G.S. is supported by a post-doctoral fellowship from the EMBO (ALTF 911-2021), T.L is a FRIA grantee of the F.R.S.-FNRS, J.K. is a Research Fellow (Aspirant) of the F.R.S.-FNRS, R.V.R. is supported by a Rubicon post-doctoral fellowship from the NWO (0191.202EN.047), G.L. is a Research Associate (Chercheur Qualifié) of the F.R.S.-FNRS.

## Author contributions

### Conceptualization

Yoann G. Santin, Géraldine Laloux

### Data Curation

Yoann G. Santin, Thomas Lamot

### Formal Analysis

Yoann G. Santin, Thomas Lamot, Jovana Kaljević, Renske van Raaphorst, Géraldine Laloux

### Funding Acquisition

Géraldine Laloux

### Investigation

Yoann G. Santin, Thomas Lamot, Jovana Kaljević

### Methodology

Yoann G. Santin, Thomas Lamot, Jovana Kaljević, Renske van Raaphorst, Géraldine Laloux

### Project Administration

Géraldine Laloux

### Resources

Géraldine Laloux

### Software

Thomas Lamot, Renske van Raaphorst

### Supervision

Géraldine Laloux

### Validation

Yoann G. Santin, Thomas Lamot, Jovana Kaljević, Renske van Raaphorst, Géraldine Laloux

### Visualization

Yoann G. Santin, Géraldine Laloux

### Writing – Original Draft Preparation

Yoann G. Santin, Géraldine Laloux

### Writing – Review &Editing

Yoann G. Santin, Thomas Lamot, Jovana Kaljević, Renske van Raaphorst, Géraldine Laloux

